# Osteochondral fluid transport in an ex vivo system

**DOI:** 10.1101/2023.10.13.562188

**Authors:** Brady David Hislop, Ara K. Mercer, Alexandria G. Whitley, Erik P. Myers, Chelsea M. Heveran, Ronald K. June

## Abstract

**Objective:** Alterations to fluid transport from bone-to-cartilage may contribute to the development of osteoarthritis. However, many questions remain about fluid transport between these tissues. The objectives of this study were to (1) test for diffusion of 3kDa molecular tracers from bone-to-cartilage and (2) assess potential differences in bone-to-cartilage fluid transport between different loading conditions.

**Design:** Osteochondral cores extracted from bovine femurs (N=8 femurs, 10 cores/femur) were subjected to either no-load (*i*.*e*., pure diffusion), pre-load only, or cyclic compression (5±2% or 10±2% strain) in a two-chamber transport system with the bone compartment filled with a 3kDa tracer. Tracer concentrations in the cartilage compartment were measured every 5 minutes for 120 minutes. Tracer concentrations were analyzed for differences in beginning, peak and equilibrium concentrations, loading effects, and time-to-peak tracer concentration.

**Results:** Peak tracer concentration in the cartilage compartment was significantly higher compared to beginning and equilibrium tracer concentrations indicating fluid transport from bone to cartilage. Cartilage-compartment tracer concentration was influenced by strain magnitude, but no time-to-peak relationship was found when comparing strain magnitudes.

**Conclusion:** This study shows that osteochondral fluid transport occurs from bone-to-cartilage with 3kDa dextran molecules. These are much larger molecules to move between bone and cartilage than previously reported. Further these results demonstrate the potential for cyclic compression to impact osteochondral fluid transport. Determining the baseline osteochondral fluid transport in healthy tissues is crucial to elucidating the potential mechanisms of progression and onset of osteoarthritis.

## Introduction

Osteoarthritis (OA) affects >500 million people each year^1^. The current standard of care for OA typically includes treatments such as pain management and exercise until an invasive total joint replacement is required. Mechanical and biological interventions that affect the trajectory of OA progression may help mitigate the physical and financial burdens of the disease. However, to design potential therapeutic interventions, a greater understanding is needed of how fluid is transported between the tissues of the synovial joint.

In the progression of osteoarthritis, structural and material changes to bone and cartilage may influence fluid transport, with repercussions to the health of the synovial joint^2-5^. In OA, the osteochondral interface (cement line, calcified cartilage, and tidemark)^6^ develops microcracks and is invaded by vasculature, leading to greater fluid transport between bone and uncalcified cartilage^7-9^. Increased fluid transport exposes cartilage to cytokines and proteases produced by bone^6, 10^. These changes occur alongside dysregulation of chondrocyte metabolism, which may contribute to other OA changes (*e*.*g*. hypertrophic chondrocytes) that drive cartilage fibrillation and degradation^11-15^. The sclerosis of subchondral bone and changes in cartilage permeability may further contribute to differences in how fluid moves within the diseased joint^16^.

While altered fluid transport has been implicated in osteoarthritis for years^16^, gaps in understanding persist about how fluid moves between healthy bone and cartilage, and how this transport depends on tissue strain. Fluid transport occurs within both cartilage and bone and contributes to mechanosensation and tissue homeostasis for both tissues^17, 18^. In addition, recent data show that small molecules (376Da, 575Da, and 1.55kDa)^19-21^ diffuse from bone to cartilage. This is significant because a permeable osteochondral interface suggests that new mechanisms for chondrocyte nutrition and cartilage-bone crosstalk (e.g., loading-induced pressure gradients that drive convection)^4^. Whether larger molecules can travel from bone to cartilage is not yet tested but is important to understand because biological molecules that would be found within bone, such as insulin (5kDa), insulin like growth factor (6.5kDa), and bone morphogenetic protein-1 (16kDa), are at least 3kDa in size. Furthermore, if the permeability changes with the progression of OA^16, 22, 23^, the disrupted crosstalk may influence OA progression^6, 24, 25^. Deformation-induced convective transport (*i*.*e*. that driven by compression induced interstitial fluid flow) may also play an important role in molecular transport from bone to cartilage. However, the effects of compression on fluid transport from bone to cartilage in healthy tissues are undetermined. Defining the relationship between fluid transport and strain is necessary for improving our understanding of bone-cartilage fluid transport in synovial joints.

In this study, we look to address several critical gaps in our understanding of fluid transport across the osteochondral interface. First, we test for diffusion of 3kDa fluorescent dextran markers across the osteochondral interface, specifically from bone-to-cartilage. Next, we look to identify potential differences in osteochondral fluid transport between different loading conditions. Finally, we test for differences between time-to-peak concentration for different loading conditions. Measured concentrations for each experimental group were compared to determine potential effects of strain magnitude and time on osteochondral fluid transport. The results provide evidence that that bone-cartilage fluid transport of uncharged molecules at least 3kDa in size most likely occurs in healthy joints and has the potential to enable crosstalk between these tissues that is not mediated through diffusion alone.

## Methods

### Bioreactor development

A two-chamber bioreactor was designed and manufactured for these studies (**Figure 1A**). A 1.5”x1.5”x1-3/16” base with a 15/16” diameter (3/8” depth), and a threaded hole on bottom for attachment to the load cell. The top chamber of the bioreactor is separated by a 1/32” rubber sheet of 1.5”x1.5”x1/32” from the base and includes four vertical through holes to secure the two chambers, as well as two horizontal through-holes (separated by 90°) to circulate fluid during experimental runs. Two identical bioreactors were manufactured using polysulfone (McMaster-Carr, Elmhurst, IL USA). The bioreactors included shims in both chambers to match fluid volumes (∼2mL). The shim in the bottom chamber (*i*.*e*., bone fluid chamber) was 3D printed with a ∼7mm hole at ∼2.4mm in depth, allowing for the press fit of the bone side of osteochondral samples. Control experiments were performed with an impermeable 8mm diameter aluminum cylinder to test for leakage between the bottom and top chambers.

**Figure 1.**
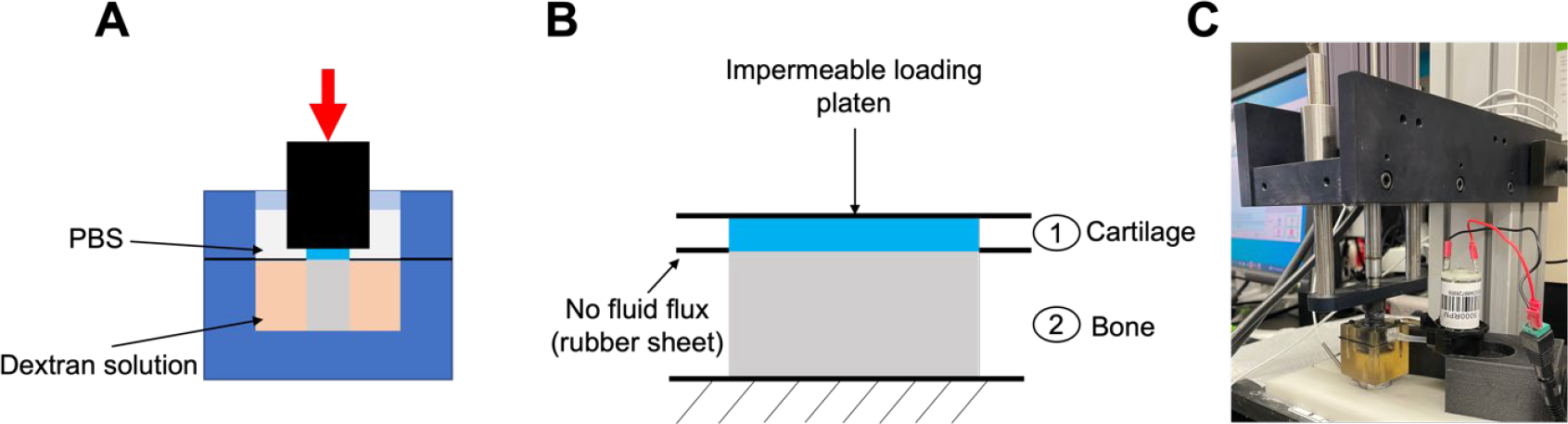
Schematics and visualization of experimental conditions for osteochondral fluid transport studies A) Cross-section of the bioreactor and loading platen, showing the cartilage fluid chamber with PBS separated from the bone fluid chamber with dextran solution by a rubber sheet B) Free body diagram showing the forces and boundary conditions acting on the osteochondral cores C) Experimental set-up for loading studies showing the peristaltic pump attached to the bioreactor circulating the fluid in the cartilage chamber.

### Bovine osteochondral core extraction

Bovine femurs were obtained from a local butcher (N=8). During sample processing the distal ends of the femurs were hydrated consistently with phosphate buffered saline (PBS). First, the femur was trimmed to create two flat surfaces directly posterior to the condyles using a band saw (Central Machinery Bench top band saw, Supplemental Figure 1). Next using an 8 mm bone coring reamer (Arthrex Osteochondral Autograft Transfer System, coring reamer) attached to a drill press (Central Machinery, 12-speed bench drill press), ten cores were drilled to a depth of ∼20 mm (n=5 per condyle). The bone coring tool was continuously hydrated with ice water to minimize the temperature increase during coring. Finally, the cores were extracted by trimming the condyles further with the band saw. Once each of the cores were removed, they were immediately placed in 2mL Eppendorf tubes with PBS and stored at -80°C until experimental day^26^.

### Osteochondral core experimental preparation

Experiments were randomized using a custom randomization script for each experimental day (Supplemental File 1), assigning the cores to an experimental group. Four experimental groups included pure-diffusion (*i*.*e*. no-loading), pre-load only, 5±2% cyclical compression, or 10±2% cyclic compression. For the groups with cyclical compression, the frequency was 1.1 Hz to simulate walking. On the day of the experiment, cores were removed from the -80°C freezer and placed in a 37°C water bath for 5 minutes. The PBS was then replaced, and the core was allowed to equilibrate for 33 minutes^26^. After equilibration, cores were trimmed to ∼12mm in height (height was not measured precisely to limit processing time), and the bone face was ground to create a perpendicular angle between the loading platen and the articular surface. A 1/32” thick rubber sheet (McMaster-Carr, 1/32”) with 3mm diameter centered hole was pulled over the core until snug around the osteochondral interface (**Figure 1A-B**), the core was then placed into the bioreactor with the bone side into the bottom transport chamber (bone fluid chamber).

### Experimental procedure

After securing the core in the bioreactor with the rubber sheet snug around the osteochondral interface, the cartilage chamber was secured to create two separate fluid chambers. 2mL of 13μM 3kDa dextran (Invitrogen, dextran, Texas Red 3000 MW, neutral charge)^27^ was added to the bone chamber through an injection port. The port was then sealed (Gorilla Tough & Clear Double Sided Adhesive Mounting Tape). A peristaltic pump (15mL/min, Gikfun 12V DC dosing pump peristaltic) was then connected to the fluid ports, and the cartilage chamber was filled with 2mL of PBS. For pure diffusion (*i*.*e*. no-load) studies two 100μL aliquots (t=0^-^) were immediately sampled from the cartilage chamber and placed in a 96-well plate (Fisherbrand™ 96-Well Polystyrene Plate). The sampled fluid was immediately replaced with 200μL of PBS and the pump was started, after which two more 100μL aliquots (t=0^+^) were sampled and again replaced with 200μL of PBS. Subsequently aliquots were sampled with PBS replacement from the cartilage chamber every 5 minutes for two hours. For pre-load only and cyclic compression groups, immediately after connecting the pump and placing 2mL PBS in the cartilage chamber two 100μL aliquots (t=0^-^) were sampled and replaced in volume. Next, a 3.5N compressive preload was applied (∼70kPa), and the position of loading platen was recorded to define the 0% strain level. The strain positions were automatically calculated using a Python script (Supplemental file 2) and input to the machine for cyclic compression studies. Immediately after pre-load two 100μL aliquots (t=0^+^) were sampled, and replaced with 200μL of PBS as the cyclic compression experiments were started. Aliquots were then sampled from the cartilage chamber every 5 minutes for two hours and replaced with 200μL of PBS each time. For pre-load only studies the loading platen was placed well above the bioreactor, and the t=0^+^ sample were taken ∼10-15s after pre-load. Between each set of aliquots, the 96-well plate was covered with aluminum foil to minimize light exposure.

### Fluorescent intensity studies

Each day a calibration curve was generated to calculate dextran concentration (1.3nM-2.8μM) from fluorescence intensity using a 96-well plate. At the end of each experiment the respective plates were analyzed with a fluorescent plate reader (BioTek, Synergy H1 Microplate reader). Excitation/emission of 595/615nm were used to measure the fluorescent intensity of each well. All concentrations were fit using a 2^nd^-order polynomial without an intercept and used to determine concentrations for their given experimental days.

### Statistical analysis

#### A. Analysis of dependence of peak intensity on experimental group

Cartilage fluid chamber concentrations at beginning (t=0^-^), peak, and equilibrium were analyzed using mixed model analysis of variance (ANOVA). The peak tracer concentration was evaluated at 15-minutes for consistency, as the time to peak concentration was not consistent. Fixed effects included strain magnitude (*i*.*e*. no load, pre-load only, 5% strain, 10% strain), and time-point (*i*.*e*. beginning, peak, equilibrium). The random effects in the model included the location on the femur where tissue was harvested, and the specimen ID. The model included the interaction term between the strain magnitude and the time-point. Measured concentrations were natural log transformed, if necessary, to meet ANOVA assumptions of residual normality and equal variance. *Post hoc* analysis was performed for significant main effects (when greater than 2 levels) or for interactions, while maintaining a family-wise error rate of 0.05 using Tukey’s multiple comparisons correction.

#### B. Analysis of dependence of peak intensity on time

Time to peak concentration was analyzed using one-way ANOVA. We considered the effect of experimental condition. All assumptions of ANOVA were verified prior to interpretation of results.

## Results

### Validation of two-fluid chamber bioreactor

Prior to performing fluorescent tracer concentration studies for the experimental conditions, we validated the baseline change in tracer concentration between the two fluid chambers using an impermeable aluminum cylinder. For all tests (n=3) we placed 13μM dextran solution in the bone fluid chamber. First, we examined the system leakage by placing a rubber sheet with no center hole and found minimal change in tracer concentration in the cartilage chamber (i.e. upper fluid chamber). Next, using the aluminum cylinder (8mm diameter) we determined the baseline tracer concentration increase in the system to be ∼5nM, indicating that detectable results would require dextran concentrations greater than this value (**Figure 2**).

**Figure 2.**
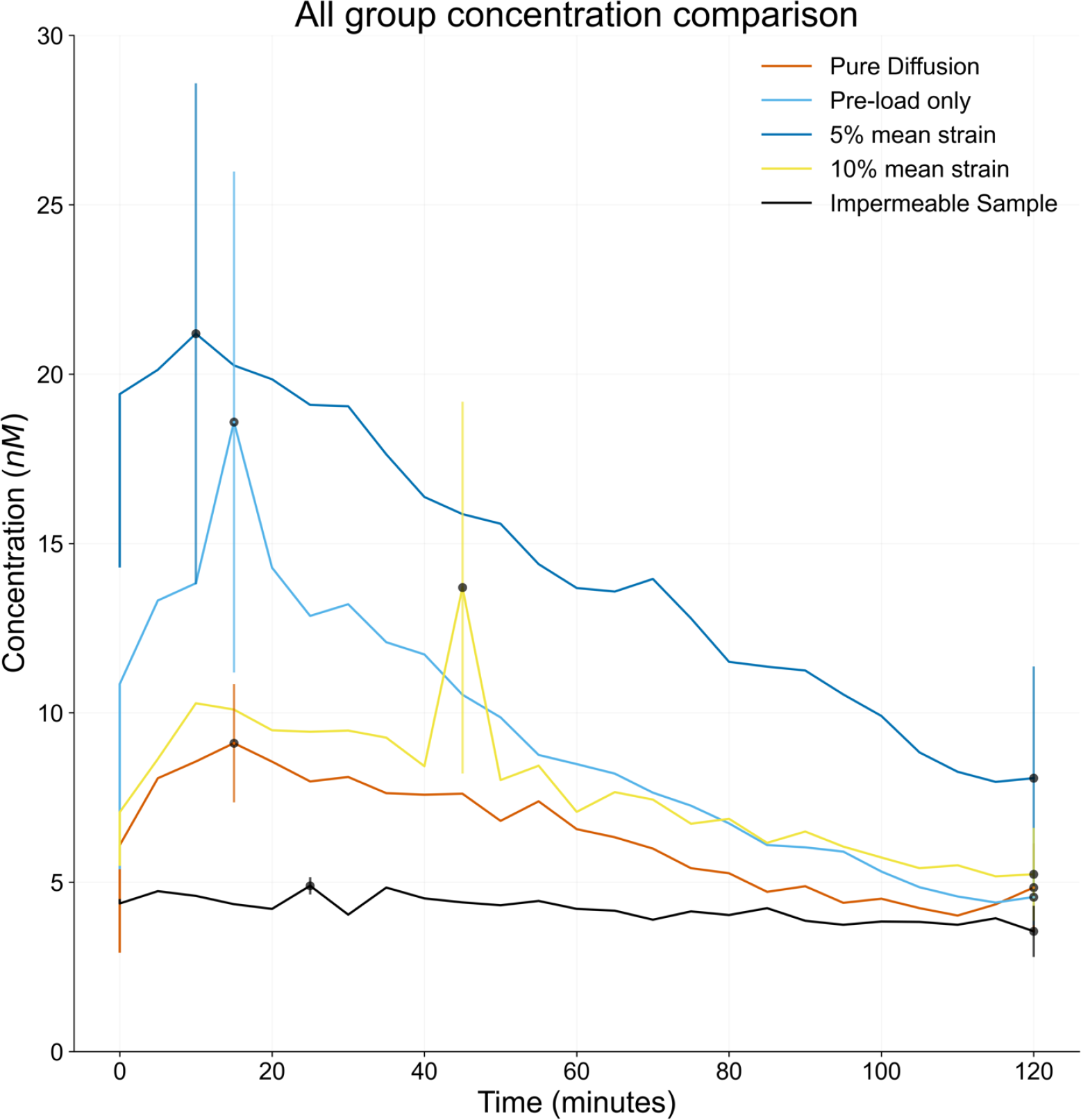
Comparison of mean tracer concentrations for experimental groups and impermeable sample, showing the dynamics of tracer concentration changes over two hours compared to the steady state impermeable tracer concentration.

**Figure 3.**
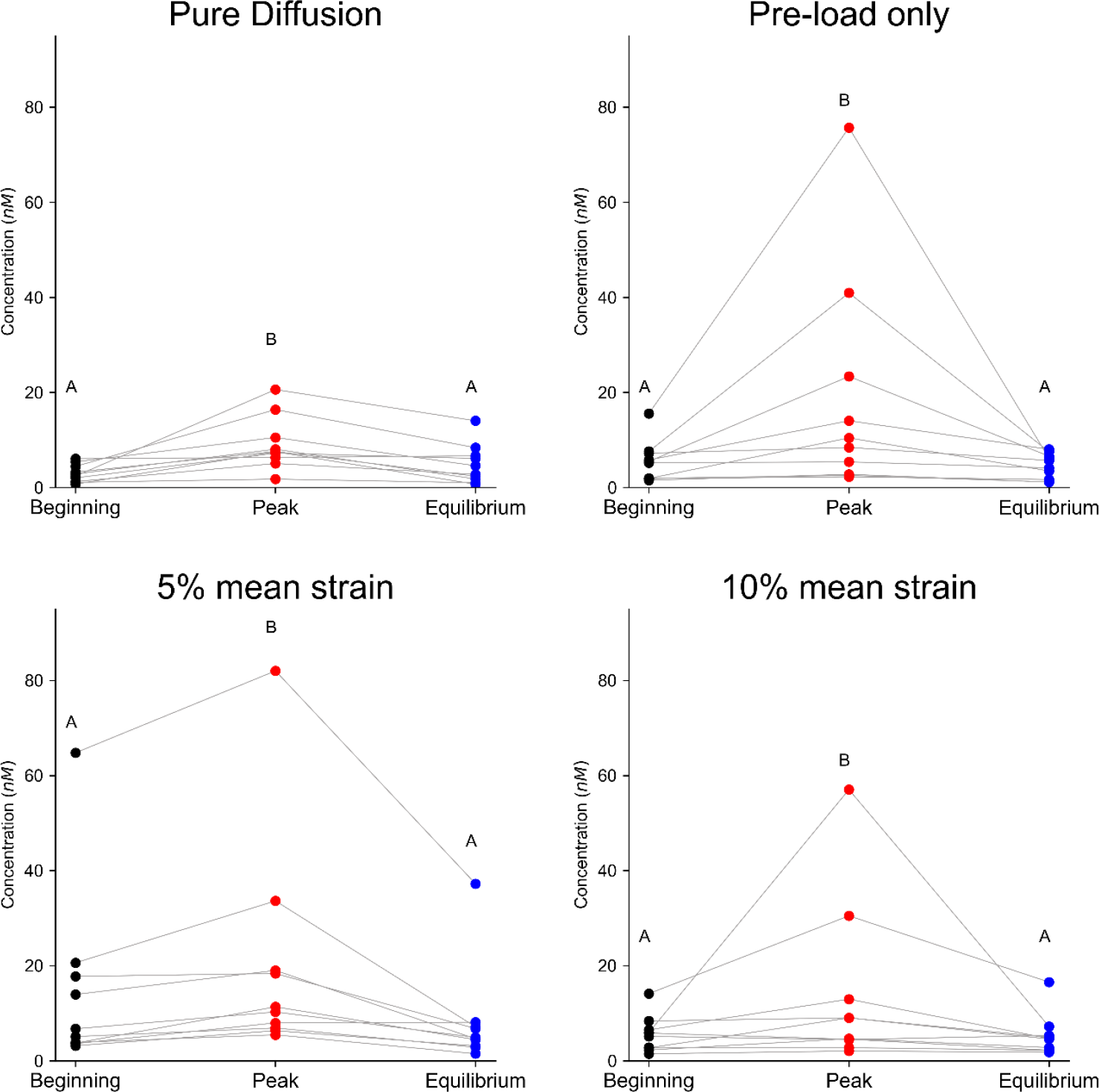
Line series plots showing changes in tracer concentration between beginning (0^-^), peak, and equilibrium (120 minutes). A) Pure-diffusion studies B) Pre-load only studies C) 5% mean strain studies D) 10% mean strain studies. Mean represented by the black crossbar, median represented by the orange crossbar in each violin plot. For each subplot the letters A and B represent group differences.

### Dextran tracer concentration changes with time and depends on strain

Tracer concentrations in the cartilage fluid chamber with significantly different between beginning, peak, and equilibrium times (p<0.001), and between strain magnitudes (p<0.001) (**Table 1**). Post hoc comparisons of time points revealed a significantly higher tracer concentration at peak compared to both beginning (p<0.001) and equilibrium (p<0.001) times, demonstrating osteochondral fluid transport from bone to cartilage. Furthermore, post hoc comparisons of strain magnitudes revealed significantly higher tracer concentrations for the 5% mean strain magnitude compared to the pure-diffusion group (p<0.001). All other post hoc comparisons were not significant. These results suggest that 5% mean strain impacts the transport of 3kDa dextran molecules from bone-to-cartilage.

**Table 1.** Summary statistics for each strain magnitude at beginning, peak and equilibrium (mean ± standard error).

| *Strain Magnitude* | *Beginning* | *Peak* | *Equilibrium<sup>3</sup>* |
| --- | --- | --- | --- |
| Impermeable | 4.47±0.50 | 4.89±0.26 | 3.55± 0.76 |
| Pure Diffusion | 2.95±0.58 | 9.11±1.75 | 4.84± 1.31 |
| Pre-load only | 5.41±1.35 | 18.58±7.40 | 4.56± 0.83 |
| 5% mean strain | 14.32±5.96 | 21.19±8.76 | 8.07± 3.30 |
| 10% mean strain | 5.52±1.18 | 13.70±5.49 | 5.23± 1.37 |

### Comparison of time to peak equilibrium reveals no effect of experimental condition

The time at which the peak tracer concentration was reached was not significantly impacted by experimental conditions (p=0.450).

## Discussion

Osteochondral fluid transport has long been neglected in the literature until the onset of OA where vascular invasion into articular cartilage is observed^8, 9, 28^. The osteochondral interface, which spans from the cement line to the tidemark is considered as an impermeable boundary preventing any molecular crosstalk between bone and cartilage^29, 30^. Thus, most studies consider synovial fluid to be the sole source of nutrients for articular cartilage. However, recent studies with small tracer molecules and the data presented here show the limitations of this prior understanding^4, 20, 23, 31^

This study provides novel data on the transport of 3kDa molecules from bone-to-cartilage (*i*.*e*. osteochondral fluid transport) in an *ex-vivo* osteochondral core system. We found that tracer concentrations changed with time, reaching a peak a few minutes after the start of the experiment. These results, together with prior studies with smaller molecules (e.g. 376Da, ∼twice the size of glucose), provide evidence that a variety of uncharged molecules can move between bone and cartilage ^20^. Bone is highly metabolically active and small molecules produced by this tissue could have roles in crosstalk with cartilage that are currently underappreciated. Future studies should consider a broader range of molecule sizes, and the effects of charged molecules on diffusion of fluid from bone-to-cartilage.

We also investigated whether fluid transport between bone and cartilage depends on the magnitude of cyclic compressive strain. Our results demonstrate that the 5% +/- 3% strain group did increase the peak tracer concentration compared with the pre-load condition. Interestingly, the other strain groups (including 10% +/-2% strain) did not differ from the pre-load. A potential explanation for this lack of influence by the larger strain could be that the cartilage was not fully unloaded (i.e., tissues were unloaded to 8% strain, not to zero strain). It is possible that pore networks of cartilage cannot fully open without greater unloading, limiting the ability of fluid to convectively and/or diffusively move into cartilage from bone. The range of 2%-5% cyclic strain in the lower strain magnitude group may better allow opening and closing of pore networks in cartilage. However, whether pore network opening in cartilage is an important regulator of convective fluid transport between bone and cartilage requires further investigation. Studies should consider a greater range of peak and unloading strains as well as loading and unloading rates.

Finally, we investigated the impacts of time to peak concentration for each strain magnitude to understand the impacts of loading on tracer concentration with respect to time. Time to peak concentration was not significantly impacted by strain magnitude. These results support our findings of convective fluid transport from bone-to-cartilage, as the independence of time to peak tracer concentration and strain magnitude removes the potential explanation of changes in diffusion properties caused by cyclic compression.

There are several limitations to this study. The temporal resolution in which aliquots were sampled every five minutes likely misses the early concentration dynamics of our system. Future studies should use faster sampling for the fluorescent measurements. Other limitations include the need to sample and replace the volume of the aliquots every five minutes with PBS likely diluting the observed tracer concentrations in the cartilage chamber. Hence, the design of a continuous fluorescent measurement system would improve temporal resolution and eliminate the issue of tracer concentration dilution. Studying the changes in tracer concentration of the cartilage fluid chamber provide evidence of bulk fluid transport from bone-to-cartilage but do not provide insight on the location from which the fluid is moving from bone-to-cartilage. Future studies should look to couple tracer concentration measurements with imaging to determine the spatial extent of the fluid movement from bone-to-cartilage. Our study focused on the study of one molecular size (3kDa), yet many other important molecules exist of varying sizes. Future studies should look to investigate a broader range of physiologically relevant molecule sizes^27^, and should investigate globular macromolecules^32^. Finally, our study does not consider the mechanical properties of the bone and/or cartilage and its impact on the bone-to-cartilage fluid transport. To understand such impacts future studies might incorporate finite element modeling to studying the mechanics of bone-to-cartilage fluid transport, or simultaneously characterize the mechanical properties of bone and cartilage looking for correlations between bone-to-cartilage molecular flux and the mechanical properties.

In conclusion, we found bone-to-cartilage fluid transport in an *ex vivo* system, suggesting that the current paradigm of an impermeable osteochondral interface should be reconsidered. The implications of increased bone-to-cartilage fluid transport in health are not well understood and need significant future investigation. By understanding the amount of fluid transport from bone-to-cartilage we can better understand the nutrient environment of articular cartilage. Overall, by establishing the baseline molecular flux from bone-to-cartilage we will improve our understanding of joint physiology which may lead us one step closer to new treatments for OA.

## Author contributions

Experimental design: BDH, RKJ; Bioreactor Development: BDH, EPM, RKJ; Data collection: BDH, AKM, AGW; Data analysis and interpretation: BDH, CMH, RKJ; Manuscript drafting: BDH, RKJ; Revision of manuscript: all authors; Approval of manuscript: all authors

## Conflict of Interest

Dr. June owns stock in Beartooth Biotech and OpenBioWorks, which were not involved in this study.

## Role of the funding source

This work was supported by grants from the National Institutes of Health (NIAMS R01AR073964 and R01AR081489) and the National Science Foundation (CMMI 1554708). This work represents the views of the authors and not necessarily those of the sponsors.

## Acknowledgements

We would like to thank Dr. Priyanka Brahmachary for procuring the bovine femurs, and Alexander Springer for helping with the initial experimentation.

## Supplemental Figures

**Supplemental Figure 1.**
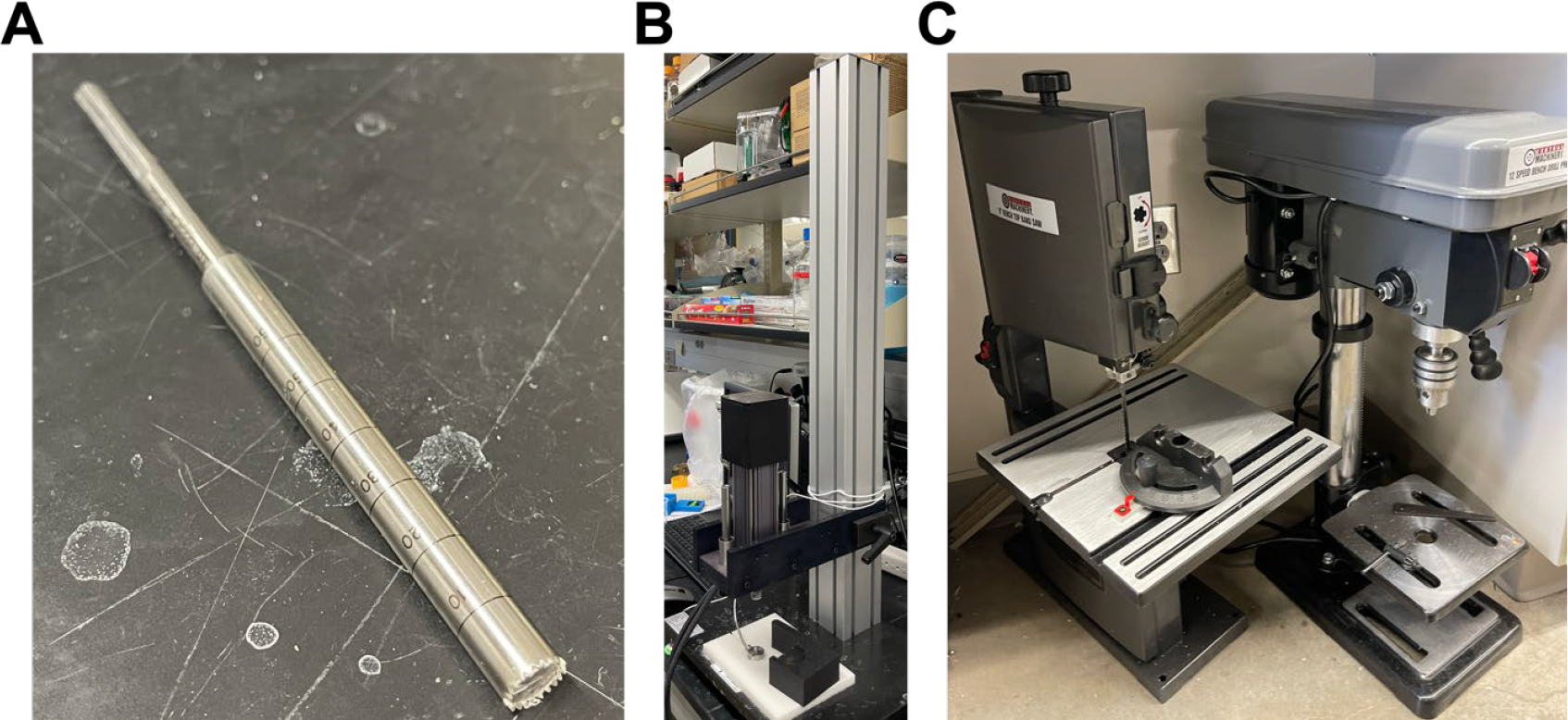
Machinery and tools used to extract cores A) Osteochondral coring reamer B) Load frame used to cyclic compression, and pre-load only studies C) Band saw (left), and drill press (right) used to extract the osteochondral cores.

## Notes

### Competing Interest Statement

The authors have declared no competing interest.

